# Single Molecule Mechanics and Kinetics of Cardiac Myosin Interacting with Regulated Thin Filaments

**DOI:** 10.1101/2023.01.09.522880

**Authors:** Sarah R. Clippinger Schulte, Brent Scott, Samantha K. Barrick, W. Tom Stump, Thomas Blackwell, Michael J. Greenberg

**Affiliations:** Department of Biochemistry and Molecular Biophysics, Washington University School of Medicine, St. Louis, MO, 63110, USA

**Keywords:** optical trapping, single molecule, myosin, thin filament

## Abstract

The cardiac cycle is a tightly regulated process wherein the heart generates force to pump blood to the body during systole and then relaxes during diastole. Disruption of this finely tuned cycle can lead to a range of diseases including cardiomyopathies and heart failure. Cardiac contraction is driven by the molecular motor myosin, which pulls regulated thin filaments in a calcium-dependent manner. In some muscle and non-muscle myosins, regulatory proteins on actin tune the kinetics, mechanics, and load dependence of the myosin working stroke; however, it is not well understood whether or how thin filament regulatory proteins tune the mechanics of the cardiac myosin motor. To address this critical gap in knowledge, we used single-molecule techniques to measure the kinetics and mechanics of the substeps of the cardiac myosin working stroke in the presence and absence of thin filament regulatory proteins. We found that regulatory proteins gate the calcium-dependent interactions between myosin and the thin filament. At physiologically relevant ATP concentrations, cardiac myosin’s mechanics and unloaded kinetics are not affected by thin filament regulatory proteins. We also measured the load-dependent kinetics of cardiac myosin at physiologically relevant ATP concentrations using an isometric optical clamp, and we found that thin filament regulatory proteins do not affect either the identity or magnitude of myosin’s primary load-dependent transition. Interestingly, at low ATP concentrations, thin filament regulatory proteins have a small effect on actomyosin dissociation kinetics, suggesting a mechanism beyond simple steric blocking. These results have important implications for both disease modeling and computational models of muscle contraction.

**Significance Statement:** Human heart contraction is powered by the molecular motor β-cardiac myosin, which pulls on thin filaments consisting of actin and the regulatory proteins troponin and tropomyosin. In some muscle and non-muscle systems, these regulatory proteins tune the kinetics, mechanics, and load dependence of the myosin working stroke. Despite having a central role in health and disease, it is not well understood whether the mechanics or kinetics of β-cardiac myosin are affected by regulatory proteins. We show that regulatory proteins do not affect the mechanics or load-dependent kinetics of the working stroke at physiologically relevant ATP concentrations; however, they can affect the kinetics at low ATP concentrations, suggesting a mechanism beyond simple steric blocking. This has important implications for modeling of cardiac physiology and diseases.

## Introduction

The human heart is finely tuned to generate the appropriate power necessary to pump blood in response to a wide range of physiological and pathological stimuli. This power output, driven by the interactions between β-cardiac myosin and the thin filament, has exquisite regulatory mechanisms to ensure that the heart generates sufficient power during systole to perfuse the body and then relaxes during diastolic filling (1,2). At the molecular scale, these regulatory mechanisms include the calcium-dependent gating of the interactions between myosin and the thin filament by the proteins troponin and tropomyosin (3), load-dependent effects on myosin kinetics (4,5), and thick filament dependent regulation (6). Dysfunction of these regulatory mechanisms can lead to cardiomyopathy, arrhythmias, and heart failure (7–10), and these regulatory mechanisms have emerged as therapeutic targets (11–13).

β-cardiac myosin (*MYH7*) is the motor that drives cardiac contraction in human ventricles, and it has different mechanochemical properties from α-cardiac myosin (*MYH6*), the motor that drives the contraction of human atrial and mouse ventricular tissues (2,14,15). Recent optical trapping experiments elucidated the mechanical and kinetic parameters that define the interactions between single β-cardiac myosin motors and actin (4,5,16,17). These studies demonstrated that β-cardiac myosin has a two substep working stroke, where the myosin generates ~6 nm total displacement. Moreover, these studies showed that mechanical forces which oppose the β-cardiac myosin working stroke slow the kinetics of ADP release from actomyosin, leading to slowed crossbridge cycling kinetics. This load-dependent slowing of cycling kinetics contributes to the force-velocity relationship in muscle, which is a critical determinant of power output (4). Importantly, this relationship can be impaired by mutations that cause disease, and there are compounds in clinical trials for cardiovascular diseases that target this relationship (16).

While previous studies have defined the mechanical and kinetic properties of β-cardiac myosin interacting with actin, myosin in the heart interacts with regulated thin filaments, which are macromolecular complexes consisting of actin and the regulatory proteins troponin and tropomyosin. It is currently not known whether these thin filament regulatory proteins affect the mechanics and/or kinetics of the β-cardiac myosin working stroke. Optical trapping experiments with other myosin isoforms demonstrated that regulatory proteins on actin can sometimes affect the mechanics and kinetics of the motor. For example, it was shown that tropomyosin Tm4.2 can increase the force sensitivity of non-muscle myosin-IIA (18). Moreover, the yeast myosin-V isoform Myo2p walks processively only in the presence of tropomyosin Tpm1p, since tropomyosin decorated actin filaments slow the rate of ADP release from the motor (19). Some studies with skeletal muscle myosin showed that thin filament regulatory proteins reduce the size of the myosin working stroke by half, potentially through disruption of the interactions between myosin’s two heads (20), while other studies showed no effect (21,22).

Defining the fundamental parameters underlying heart contraction is critical for understanding and modeling cardiac physiology and disease. It is essential to determine how regulated thin filaments, not just actin, tune the mechanics, kinetics, and/or load-dependent properties of β-cardiac myosin. Therefore, we used high-resolution optical trapping methods to study these properties in the presence of fully regulated thin filaments.

## Results

### Reconstitution of regulated thin filaments

Cardiac actin and β-cardiac myosin were tissue purified from porcine ventricles. Porcine cardiac actin is identical to human actin, and porcine β-cardiac myosin is 97% identical to human β-cardiac myosin, with biochemical and biophysical properties that are indistinguishable from the human isoform (4,5,14,17,23–25). Human troponin and tropomyosin were expressed recombinantly and reconstituted with purified actin to form regulated thin filaments. In vitro motility assays demonstrate that these thin filaments are functional, with no movement at low calcium (pCa 9) and robust movement at high calcium (pCa 4) (**Fig. 1a**).

**Figure 1:**
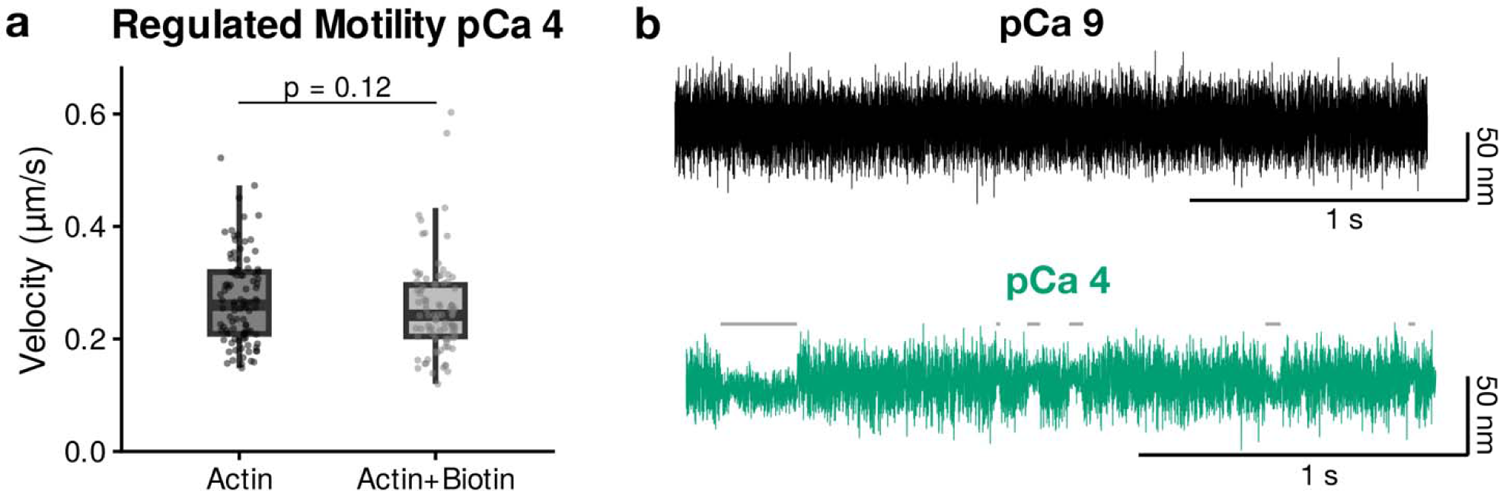
Reconstitution of thin filaments using biotinylated actin does not affect thin filament regulation. **a.** In vitro motility assay boxplots showing the speed of regulated thin filament translocation with and without biotin-labeled actin at pCa 4. Individual points show the velocity of >25 thin filaments measured across 3 separate experiments. The thick middle line of the boxplot shows the median value, the top/bottom of the box are the 1^st^ and 3^rd^ quartiles, and the whiskers extend to 1.5 times the interquartile range. There is no difference in speed with or without biotinylated actin (P = 0.12, Mann-Whitney test). No movement was seen at pCa 9 with or without biotinylated actin. **b.** Representative optical trapping traces of regulated thin filaments containing 10% biotinylated actin conducted at low (pCa 9, orange) and high calcium (pCa 4, blue). At low calcium, binding events were very rare and at pCa 4 binding events were frequent (grey lines denote binding interactions detected by the analysis program), demonstrating regulation.

For the optical trapping experiments, it is necessary to attach regulated thin filaments to polystyrene beads via a linkage. In these experiments, we mixed 10% biotinylated G-actin with unlabeled G-actin during polymerization and then used the biotinylated linkage to attach the reconstituted thin filament to streptavidin-coated beads (4). To ensure that the biotinylated actin does not interfere with thin filament regulation under the same fully activating or inhibiting conditions used in the trapping assays, we performed in vitro thin filament gliding assays at both low (pCa 9) and high (pCa 4) calcium (26). Calcium-based regulation of these thin filaments was observed, with movement at pCa 4 and no movement at pCa 9. There was no significant difference in the average speed of thin filament translocation at pCa 4 between regulated thin filaments made from unlabeled actin (0.26 ± 0.08 μm/s) and those including 10% biotinylated actin (0.25 ± 0.08 μm/s) (P = 0.12) (**Fig. 1a**).

Next, we tested whether we could observe calcium-dependent thin filament regulation in our optical trapping assay. We used the three-bead assay pioneered by the Spudich lab (27), in which two optically trapped beads are attached to a single actin filament and then lowered onto a surface-bound bead that is sparsely coated with myosin. Using reconstituted regulated thin filaments attached to streptavidin-coated polystyrene beads, we examined binding interactions between myosin and the regulated thin filaments. At 1 μM ATP, we could clearly resolve binding interactions between regulated thin filaments and myosin at high calcium; however, these binding interactions were exceedingly rare at low calcium (**Fig. 1b**). Taken together, our results demonstrate that we are able to reconstitute functional regulated thin filaments for optical trapping assays.

### Regulatory proteins do not affect the mechanics of the myosin working stroke at low ATP

To examine the effects of regulatory proteins on the mechanics of the β-cardiac myosin working stroke, we used the three-bead optical trapping assay. Experiments were conducted at 1 μM ATP and high calcium (pCa 4) to facilitate observation of substeps of the myosin working stroke (28). Experiments were conducted in both the presence and absence of regulatory proteins. Myosin binding to actin causes a reduction in the variance of the bead position, and individual interactions between myosin and thin filaments could clearly be resolved (**Figs. 2a-b**). Binding interactions were resolved using a covariance threshold (29). Cumulative distributions measuring the total size of the myosin working stroke show that data are well described by a single Gaussian function (**Fig. 2e;** P = 0.09 for unregulated and P = 0.25 for regulated filaments). The total size of the β-cardiac myosin working stroke measured with unregulated actin (5.3 ± 8.6 nm; n = 364 binding events) was consistent with previous measurements (4,17,24,29) and not significantly different from the size of the working stroke measured with regulated thin filaments (4.9 ± 9.1 nm; n = 491 binding events; P = 0.57).

**Figure 2:**
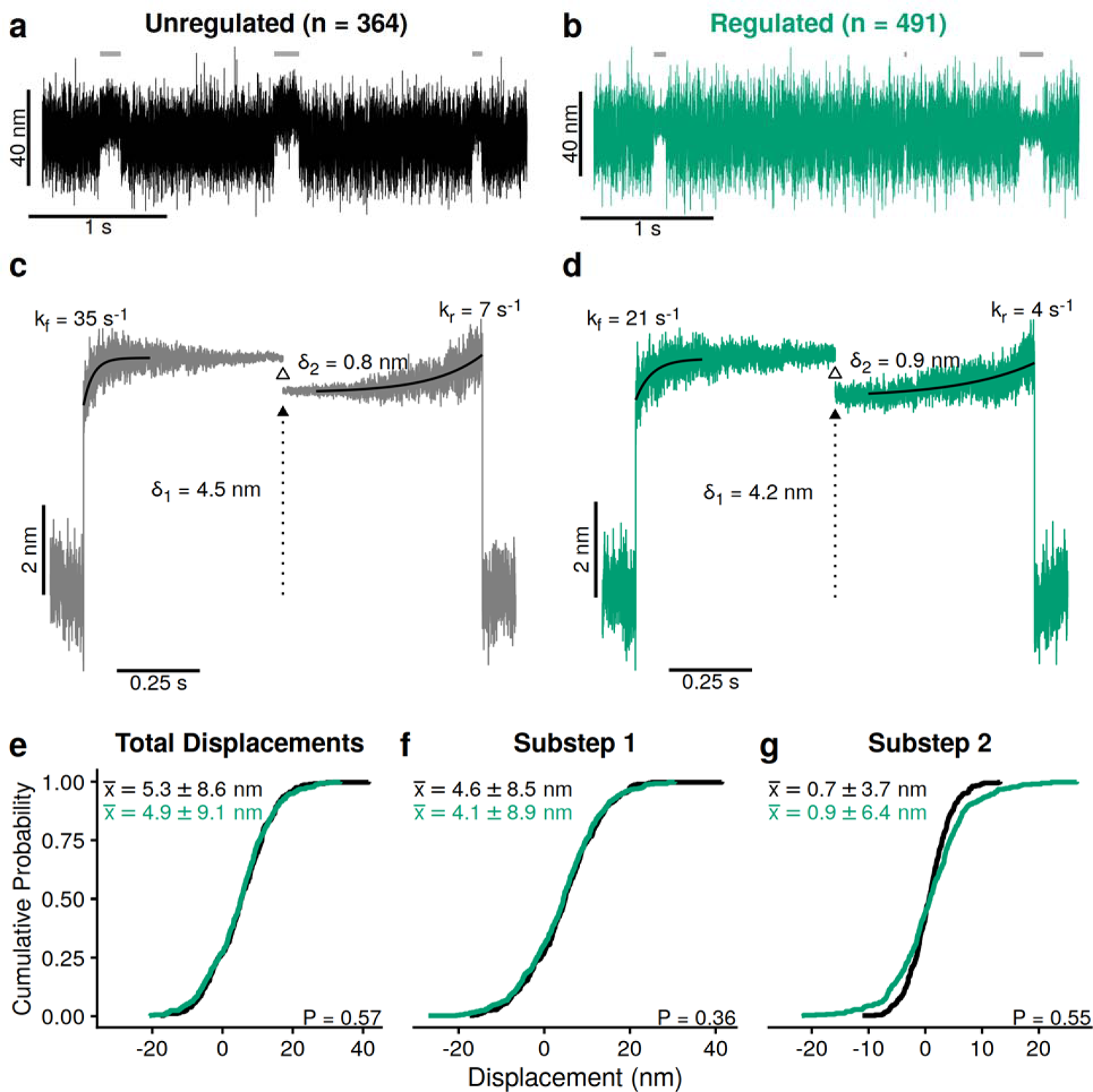
Optical trapping experiments at 1 μM ATP reveal no changes in working stroke mechanics with regulated thin filaments. Data are from 364 interactions detected from 10 myosin molecules for the unregulated condition and 491 interactions from 9 myosin molecules for the regulated condition. **a-b.** Representative optical trapping data for unregulated (black) and regulated (green) thin filaments. Grey lines indicate actomyosin interactions identified by automated event detection (see **Methods** for details). **c-d.** Ensemble averages of the working stroke for myosin interacting with unregulated (left, black) and regulated (right, green) thin filaments. The rates of the exponential functions fit to the time forward (k_f_) and time reverse (k_r_) averages are shown along with the magnitudes of the two substeps. **e-g.** Cumulative distributions derived from individual binding interactions for the (**e**) total displacement, (**f**) first substep, and (**g**) second substep. Each plot reports the sample mean, standard deviation, and P-values from t-tests.

Previous studies have shown that β-cardiac myosin accomplishes its working stroke in two substeps, with the first substep associated with phosphate release and the second associated with ADP release (4,17,25,29). To better understand the effects of regulatory proteins on the mechanics of substeps of the β-cardiac myosin working stroke, we used ensemble averaging of individual binding interactions, which enables the detection of subtle substeps that are typically obscured by Brownian motion (4,30,31). Binding interactions were synchronized either upon the initiation of binding (time forward averages) or the termination of binding (time reverse averages) using a changepoint algorithm and then averaged as previously described (29). The difference between the time forward and time reverse averages is indicative of a two-substep working stroke, and we see a two-substep working stroke for both the regulated and unregulated thin filaments (**Figs. 2c-d**), consistent with previous studies of unregulated thin filaments (4,25,29). The displacement of the time forward and time reverse averages at detachment gives the size of the total working stroke (29). The difference in displacement between the end of the time forward averages and the beginning of the time reverse averages gives the size of the second substep of the working stroke, and the difference between the total working stroke and the second substep gives the size of the first substep. Here, we constructed cumulative distributions of substep displacements from our individual binding interactions, and the data follow the expected shape of a cumulative distribution for a single Gaussian function (**Figs. 2f-g**). We did not observe a statistically significant difference in the size of either the first (P = 0.36) or second (P = 0.55) substep of the working stroke in the presence or absence of regulatory proteins.

### Regulatory proteins affect kinetics of detachment at non-physiological, low ATP concentrations

Optical trapping experiments collected at 1 μM ATP and high calcium provide information not only on the mechanics of the working stroke but also the kinetics. The distribution of attachment durations can be used to calculate the actomyosin dissociation rate. Cumulative distributions of binding interaction durations were well fitted by single exponential functions to yield the actomyosin detachment rate (**Fig. 3a**). Interestingly, we observe that the detachment rate measured in the presence of regulatory proteins (3.9 (−0.3/+0.4) s^-1^) was slightly slower than the rate measured without regulatory proteins (5.5 ± 0.4 s^-1^) (P = 0.002). This is consistent with previous studies of skeletal muscle myosin conducted at low ATP concentrations which show slower detachment rates in the presence of regulatory proteins (20).

**Figure 3:**
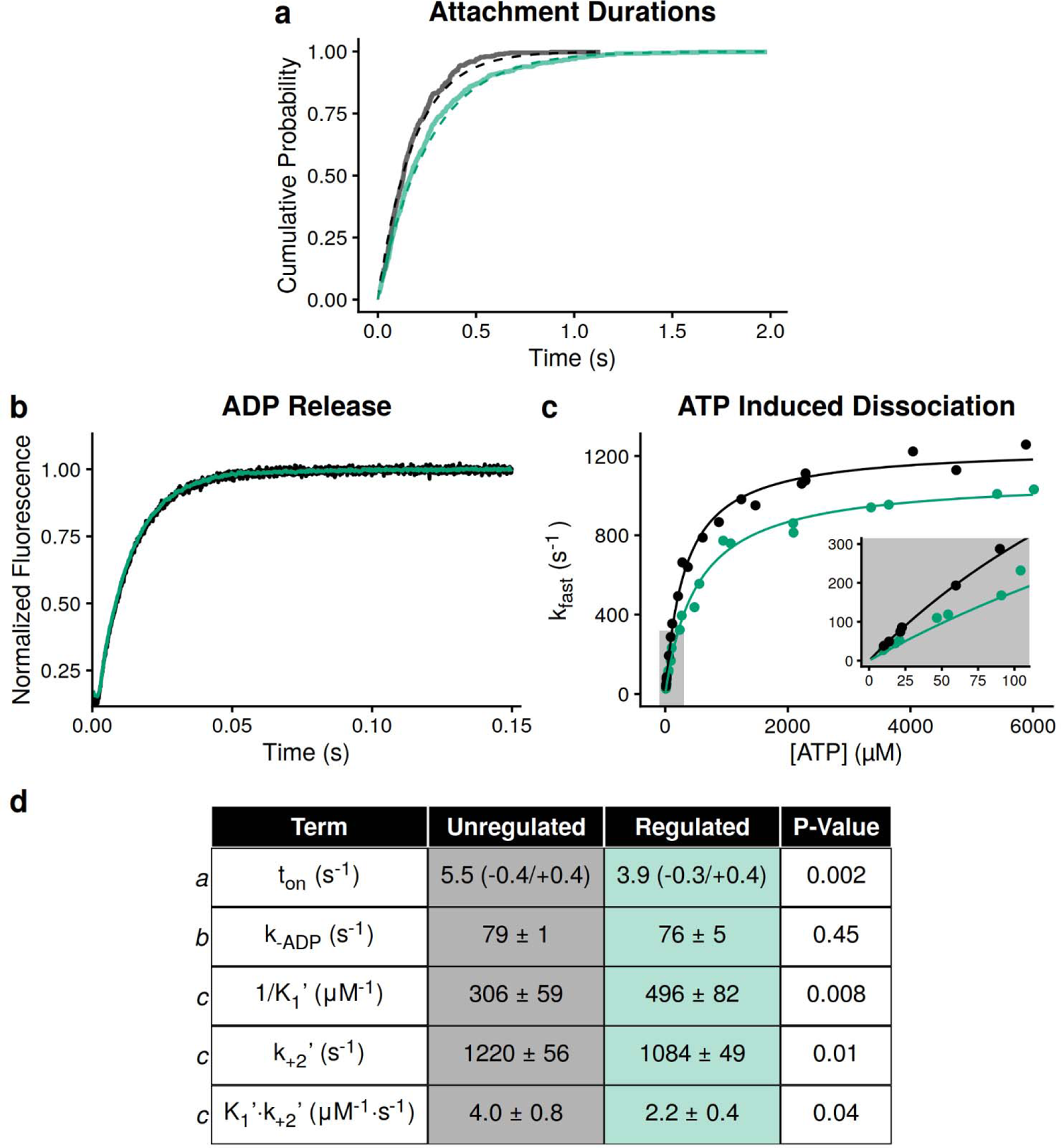
Regulatory proteins affect kinetics at low ATP due to slowing of ATP-induced dissociation. **a.** Individual actomyosin attachment durations obtained in the optical trap are plotted as cumulative distributions for unregulated (black) and regulated (green) thin filaments. Distributions are fit with single exponential functions (dashed lines) to obtain the actomyosin detachment rate. Regulatory proteins significantly slow the rate of detachment (see table in **d**; value is the fitted rate, error is a bootstrapped 95% confidence interval, and P-value is calculated as described in the **Methods**). **b.** Representative fluorescence transients from stopped-flow experiments measuring the rate of ADP release from actomyosin. Fits show single exponential fits to the average of 5 transients. There is no difference in the rate of ADP release measured using regulated or unregulated thin filaments (see table in **d**). **c.** Fast phase of ATP-induced dissociation of myosin from regulated and unregulated thin filaments (see **Methods** for details). Each point represents the average of 5 technical repeats collected on one day. 3 separate experimental days were used. Solid lines show fitting of Eq. 2. **d.** Table of parameters obtained for all kinetic measurements. The row letters indicate from which figure panel the values are derived. For the stopped-flow transient kinetics (marked “b” or “c”), the reported values are the average and standard deviations of 3 separate experimental days and P-values were calculated using a Student’s t-test.

The assertion that dissociation of regulated thin filaments from myosin is slowed at low ATP is further supported by kinetic information extracted from the ensemble averages. The time reverse averages give the rate of transitioning from the second substep to the detached state, and previous studies have shown that this transition is related to the biochemical rate of ATP-induced actomyosin dissociation (30,31). The rates of the time reverse ensemble averages measured here at 1 μM ATP are well fitted by single exponential functions (**Fig. 2c-d**). The rate of this transition measured in the presence of regulatory proteins (4 s^-1^) is slower than the rate measured in the absence of regulatory proteins (7 s^-1^). These rates are consistent with the detachment rates measured from the cumulative distributions of attachment durations, suggesting that this transition limits the actomyosin detachment rate at low ATP (**Fig. 3a**). Consistent with the notion that this transition limits detachment, the rates of the time forward averages collected at low ATP, which report the rate of transitioning from the first substep to the second (**Figs. 2c-d)**, are much faster than the rates of the time reversed averages (**Fig. 2c-d**) or the detachment rates (**Fig. 3a**). Taken together, our trapping data strongly suggest that regulatory proteins slow the rate of ATP-induced dissociation at low ATP concentrations.

To further investigate the basis for the slowed detachment rate in the presence of regulatory proteins observed in the optical trap at low ATP concentrations, we measured the rates of ATP-induced actomyosin dissociation and ADP release using stopped-flow techniques (**Figs. 3b and 3c**). It has previously been shown that at low concentrations of ATP, the rate of actomyosin dissociation is limited by the rate of ATP-induced actomyosin dissociation, while at physiologically relevant ATP concentrations, detachment is limited by the rate of ADP release (14,32,33). We found that the rates of ADP release in the presence of regulatory proteins (76 ± 5 s^-1^) and in their absence (78 ± 1 s^-1^) were not significantly different from each other (P = 0.45), and the measured values are consistent with previous studies (4,9,10,14,34). We also measured the rate of ATP-induced actomyosin dissociation. Consistent with previous studies, traces were well fitted by two exponential functions, where the rate of the fast phase (**Fig. 3c**) reports the rate of ATP-induced actomyosin dissociation, and the rate of the slow phase (**Supplemental Fig. 1**) measures the rate of a slow ATP-independent isomerization (see **Methods** for details) (35). The observed fast rate of ATP-induced dissociation was modeled as the formation of a rapid-equilibrium collision complex (K_1_’) followed by an irreversible isomerization and rapid detachment (k_+2_’) (see **Methods** for details). At saturating ATP, for both the regulated and unregulated thin filaments, the rates of ADP release were much slower than the rate of ATP-induced dissociation (**Fig. 3d**), consistent with ADP release limiting detachment at saturating ATP concentrations. At low ATP concentrations, the rate of ATP binding is given by the second-order ATP binding rate (K_1_’*k_+2_’) (**Figs. 3c inset, and 3d**). Consistent with the optical trapping measurements at low ATP (**Figs. 2c, 2d and 3a**), the second order rate of ATP binding is slower for regulated filaments (2.2 ± 0.4 μM^-1^ s^-1^) compared to unregulated filaments (4.0 ± 0.8 μM^-1^ s^-1^, P = 0.04). Taken together, these data demonstrate that regulatory proteins slow the rate of ATP-induced actomyosin dissociation at low ATP concentrations; however, at physiologically relevant ATP concentrations, the rate of ADP release limits dissociation, and this rate is not affected by regulatory proteins.

### Regulatory proteins do not affect the loaded kinetics of cardiac myosin at saturating ATP

To examine the effects of load on myosin’s mechanics, we used an isometric optical clamp, where one optically trapped bead is assigned to be the transducer bead and the other is assigned as the motor bead (4, 36). The position of the transducer bead is measured, and the motor bead is actively moved by a feedback loop to keep the transducer bead in the center of the optical trap. When myosin binds to the actin and pulls, both beads are pulled from the centers of their traps. The motor bead is then pulled to return the transducer bead back to its original position, exerting a load on the myosin (**Fig. 4a**). The attachment duration and the load on the myosin are measured. Using this technique, a range of forces can be measured. Importantly, these experiments were conducted at physiologically relevant saturating ATP concentrations (1 mM) to enable the probing of mechanical states that are populated in the healthy myocardium (4,16).

**Figure 4:**
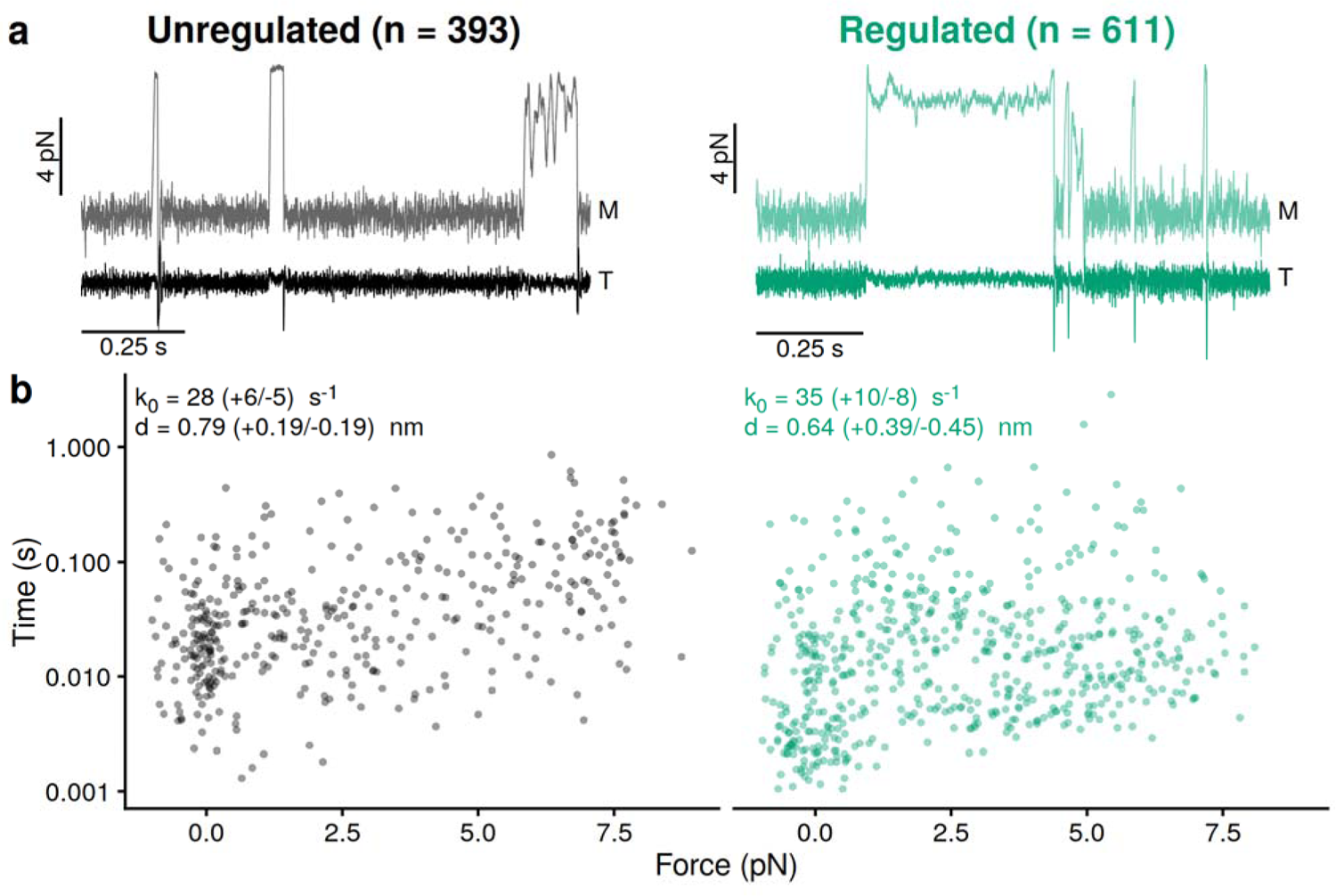
Regulatory proteins do not affect myosin’s load-dependent kinetics at physiological ATP. **a.** Representative traces collected using the isometric optical clamp. The motor bead (“M”) is moved to keep the transducer bead (“T”) at an isometric position. During a binding interaction, the average force and the attachment duration were recorded. **b.** Attachment durations as a function of force measured for unregulated (black) and regulated (green) thin filaments. Each point represents a single binding interaction. 393 binding interactions were observed from 10 myosin molecules for the unregulated condition and 611 binding interactions were observed from 20 myosin molecules for the regulated condition. Data were fitted with the Bell equation using maximum likelihood estimation to obtain k_0_ and d. Error bars are the 95% confidence intervals obtained from 1000 rounds of bootstrapping simulations. These parameters were not significantly different in the presence of regulatory proteins (P = 0.16 for k_0_ and P = 0.49 for d; see **Methods** for details on statistics and fitting).

Using the isometric optical clamp, we obtain a scatter plot where each point represents the attachment duration and average force of a single binding interaction. The distribution of attachment durations at a given force are exponentially distributed (4,37) (**Fig. 4b**). As can be seen, the attachment duration increases gradually as force is increased, consistent with force-induced slowing of actomyosin dissociation. These data were fitted with the Bell Equation (38) using maximum likelihood estimation as previously described (4,37,39) to obtain the force-dependent detachment rate:

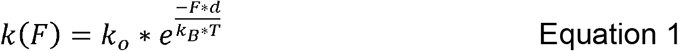

where F is the force, k_o_ is the rate of the primary force-sensitive transition in the absence of force, d is the distance to the transition state (i.e., the force sensitivity), and k_B_*T is the thermal energy, which has a value of 4.11 pN*nm at room temperature.

In the absence of regulatory proteins, we found that the rate of the primary force-sensitive transition, k_o_, is 28 (−5/+7) s^-1^. This is consistent with previous measurements in the optical trap (4,17,24,25,29) and the rate of the time forward ensemble average (35 s^-1^ **Fig. 2c**). The distance to the transition state, d, is 0.79 (−0.17/+0.19) nm, consistent with previous measurements (4,17,24,25,29). Consistent with previous work, these data demonstrate that for unregulated thin filaments, the detachment rate at saturating ATP is limited by the force-dependent rate of ADP release (4,17,24,25,29). In the presence of regulatory proteins (**Fig. 4b**), the rate of the primary force-sensitive transition, k_0_, is 35 (−8/+10) s^-1^, and the distance to the transition state, d, is 0.64 (± 0.4) nm. These values are not significantly different when comparing regulated and unregulated thin filaments (P = 0.16 and P = 0.49 for k_0_ and d, respectively). Taken together, we do not detect any differences in the force-dependent kinetics of cardiac myosin in the presence or absence of regulatory proteins at physiologically relevant saturating ATP concentrations.

## Discussion

We used high-resolution optical trapping techniques to examine the mechanics and kinetics of the β-cardiac myosin working stroke in both the presence and absence of regulatory proteins. We found that regulatory proteins do not tune the mechanics of the working stroke or the load-dependent kinetics of β-cardiac myosin contraction at physiologically relevant ATP concentrations; however, we did observe slight differences in the kinetics of actomyosin dissociation in the presence of regulatory proteins at low ATP, suggesting that these regulatory proteins have subtle effects beyond just sterically blocking the interactions between myosin and the thin filament.

### The molecular role of regulatory proteins in cardiac muscle

In the healthy heart, the ATP concentration is ~8 mM (40,41), and even in heart failure, the ATP concentration remains in the millimolar range (1-4 mM) (42–44). Here, we found that at physiologically relevant millimolar ATP concentrations, regulatory proteins have no appreciable effect on the mechanics or kinetics of the β-cardiac myosin working stroke or the kinetics of actomyosin dissociation (**Fig. 4b**). Consistent with previous studies (7,9,10,45), our stopped-flow measurements show that at physiologically relevant ATP concentrations, the rate of ADP release is much slower than the rate of ATP-induced actomyosin dissociation (**Figs. 3b-c**), and therefore the muscle shortening speed will be limited by the rate of ADP release. We did not observe any differences in the rates of ADP release (**Fig. 3b**) or the rate of actomyosin dissociation measured in the optical trap at saturating ATP (k_0_, **Fig. 4b**) in the presence of regulatory proteins. Although we observed subtle changes in the second-order rate of ATP-induced dissociation between regulated and unregulated filaments (**Fig. 3d**), these changes are irrelevant at [ATP] > 50 μM where the rate of ADP release limits detachment. Therefore, in the functioning heart, regulatory proteins do not modulate the kinetics of the myosin working stroke. Moreover, our data show that the load-dependent rate of ADP release limits actomyosin dissociation, and this is unchanged by regulatory proteins (**Figs. 3b-c and 4**). Finally, we show that the mechanics of the working stroke are unchanged by regulatory proteins (**Fig. 2**). Taken together, our data are consistent with tropomyosin primarily serving a role in sterically blocking the calcium-dependent interactions between cardiac myosin and the thin filament under working conditions in the heart (46,47).

Although not relevant to working conditions in the heart, our results reveal that regulatory proteins modulate the kinetics of ATP-induced actomyosin dissociation at low ATP concentrations. To measure the substeps of the working stroke, we conducted optical trapping experiments at very low ATP concentrations not experienced in the cell, since this slows mechanical transitions that are too fast to resolve at physiologically relevant ATP (28). While our results show that regulatory proteins do not change the size of the working stroke or the coupling between kinetics and mechanics (**Fig. 2**), they also reveal that regulatory proteins slow the dissociation of actomyosin at very low ATP concentrations (**Fig. 3a**). The time reverse ensemble averages demonstrate that actomyosin dissociation is limited by the transition from the second substep to the detached state (**Figs. 2c-d**), which is similar to the second-order rate of ATP-induced actomyosin dissociation measured using stopped flow techniques (**Fig. 3b**) (29–31). All of these rates (actomyosin detachment rate (**Fig. 3a**), time reverse ensemble average rate (**Fig. 2c-d**), and ATP-induced dissociation rate measured in the stopped flow (**Fig. 3b**)) are similar, consistent with the rate of ATP-induced dissociation limiting detachment at low ATP. Moreover, for each of these measurements, the rates are significantly slower in the presence of regulatory proteins than in the absence (**Fig. 3d**). Taken together, these data demonstrate that at low ATP concentrations, regulatory proteins slow actomyosin detachment due to slowed ATP-induced dissociation.

While not relevant to physiology, the observation that regulatory proteins can tune ATP-induced dissociation has interesting implications. The tuning of detachment kinetics by regulatory proteins cannot be fully explained by a simple steric blocking mechanism, where tropomyosin only blocks the interactions between myosin and the thin filament (46). The exact mechanism of this kinetic tuning is not known; however, it has been proposed that tropomyosin can interact directly with myosin (48), and domain-specific deletion studies of tropomyosin have demonstrated that myosin can modulate tropomyosin binding to the thin filament (49,50). Moreover, recent structural studies of muscle proteins have demonstrated specific interactions between loop-4 of myosin and tropomyosin (51,52). It is possible that these interactions are allosterically coupled to changes in the nucleotide binding pocket of myosin, given the complex allosteric networks between the nucleotide binding site and the actin binding domain (53). Moreover, our results are consistent with previous optical trapping work examining skeletal muscle myosin at low ATP concentrations, which demonstrated that regulatory proteins slow the rate of actomyosin detachment (20). That being said, our data collected at saturating ATP concentrations demonstrate that kinetic tuning by regulatory proteins is likely not relevant under working conditions in cardiac muscle, and the effects of regulatory proteins in cardiac muscle are well described by a steric blocking model.

### The effects of regulatory proteins on actomyosin appear to be isoform specific

Here, we saw that at physiologically relevant ATP concentrations, tropomyosin and troponin do not appreciably affect the β-cardiac myosin working stroke. Interestingly, the effects of regulatory proteins on myosin appear to depend on the specific protein isoforms used. There are many distinct tropomyosin isoforms expressed in eukaryotic cells, with tissue-specific expression patterns, and even within the same cell, different tropomyosin isoforms localize to different subcellular actin pools (54). Moreover, while all myosin isoforms can associate with actin, they preferentially interact with certain actin structures in an isoform-specific manner (55). The biophysical role of tropomyosin appears to vary with both the myosin and tropomyosin isoform, where some tropomyosin isoforms inhibit the interactions of certain myosin isoforms with actin (55,56), while others promote these interactions (19,57). This isoform specificity has been proposed to help localize specialized motors to specific regions of the cell (58).

The molecular roles of tropomyosins on myosin motor function can extend beyond steric effects. For example, decoration of actin with Tpm1p slows the rate of ADP release from the myosin-V isoform Myo2p, and this slowing of ADP release kinetics enables Myo2p to processively walk on tropomyosin decorated actin filaments, something it will not do on bare actin (19). Even within the family of myosin-II motors, there is evidence that regulatory proteins could tune the load-dependent kinetics of myosin in an isoform specific manner. For example, non-muscle myosin-IIA’s force sensitivity is increased by tropomyosin Tm4.2 (18). In the case of skeletal muscle myosin, some studies suggested that tropomyosin decreases the step size of myosin by inhibiting the ability of both myosin heads to bind to the thin filament (20), while other studies have not seen this effect (21,22). The working stroke size that we measured for β-cardiac myosin in the presence of regulatory proteins is indistinguishable from that measured in the absence of regulatory proteins with both one-headed and two-headed myosin constructs (4,17,24,25), so we do not believe that our inability to observe a difference in mechanics is related to the two-headed nature of the construct used here (59).

### Conclusions

Our results clearly demonstrate that under physiologically relevant ATP concentrations, regulatory proteins do not cause appreciable changes in the mechanics or kinetics of the β-cardiac myosin working stroke; however, they can tune myosin’s kinetics at low ATP concentrations, suggesting effects beyond a simple steric blocking mechanism. This has important implications for both our understanding of the mechanism of muscle regulation and mathematical modeling of muscle contraction, which relies on accurate, sensitive measurements of these parameters.

## Materials and Methods

### Protein expression and purification

Cardiac actin and myosin were purified from cryoground porcine ventricles as previously described (10). Human troponin and tropomyosin were expressed recombinantly in E. coli, purified, and complexed as described previously (10). Myosin subfragment-1 (S1) for spectroscopic measurement was prepared by limited proteolysis using chymotrypsin as previously described, and N-(1-Pyrene)Iodoacetamide-labeled actin was prepared as previously described (10). Protein concentrations were determined spectroscopically. For all experiments, at least 2 separate protein preparations were used.

### Stopped-flow measurements of ADP release and ATP-induced actomyosin dissociation

Stopped-flow experiments were performed in a SX-20 apparatus (Applied Photophysics). All experiments were conducted at 20°C in high calcium buffer (pCa 4) containing 25 mM KCl, 5 mM free MgCl_2_, 60 mM MOPS pH 7.0, 2 mM EGTA, 1 mM DTT, and 2.15 mM CaCl_2_, where the concentration of free calcium was calculated using MaxChelator (60).

ADP release experiments were performed as described previously (7,45,61). Briefly, 1 μM phalloidin-stabilized, pyrene-labeled actin, 1.5 μM tropomyosin (when appropriate), 1.5 μM troponin (when appropriate), 1 μM S1 myosin, and 100 μM Mg*ADP were rapidly mixed with 5 mM Mg*ATP. This caused an increase in fluorescence that was well fit by a single exponential function, where the rate equals the rate of ADP release (35). Each experiment consisted of 5 technically repeated measurements, and the 3 experiments were used to calculate the mean and standard deviation. A two-tailed Student’s t-test was used for statistical testing.

The rate of ATP-induced actomyosin dissociation was measured as previously described (61). Briefly, 1 μM phalloidin-stabilized, pyrene-labeled actin, 1.5 μM tropomyosin (when appropriate), 1.5 μM troponin (when appropriate), 1 μM S1 myosin, and 0.04 U/mL apyrase were rapidly mixed with varying concentrations of Mg*ATP. The resultant fluorescence transients were best fit by the sum of two exponential functions, as previously described (35). The amplitude of the fast phase was fixed to prevent artifacts due to the dead time of the instrument. As has been shown before, the rates of the fast and slow phases were hyperbolically related to the concentration of ATP. Data were interpreted according to the scheme (35):

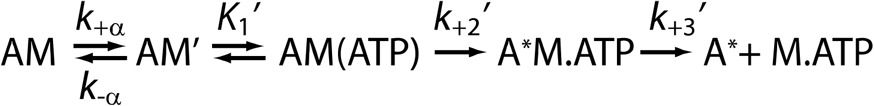

The fast phase of ATP-induced dissociation was modelled as the formation of a collision complex between actomyosin and ATP that is in rapid equilibrium (K_1_’) followed by an irreversible isomerization and rapid dissociation (k_+2_’). The rate of the fast phase, k_fast_, is hyperbolically related to the ATP concentration by:

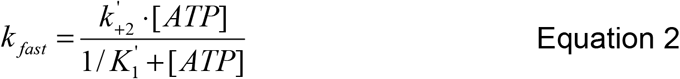

At low [ATP], the second order rate of ATP-induced dissociation is given by K_1_’ * k_+2_’. The concentration of ATP was measured spectroscopically for all experiments. The rate of ATP-induced dissociation was measured over a full range of concentrations 3 times, and each time, the fitted parameters were extracted. Reported values are the average of these 3 trials and the error is the standard deviation. Statistical testing was performed using a 2-tailed Student’s t-test.

### In vitro motility assays

*In vitro* motility assays were performed as described (10). For the experiments using biotinylated actin, all experimental protocols were identical except 10% biotin-actin (Cytoskeleton) was added to 2 μM G-actin and stabilized with tetramethylrhodamine isothiocyanate-labeled phalloidin in KMg25 buffer (60 mM MOPS pH 7.0, 25 mM KCl, 2 mM EGTA, 4 mM MgCl_2_, and 1 mM DTT). The concentration of free calcium was calculated using MaxChelator (60). Reported values are the average and standard deviations of the speeds from at least 3 separate days of experiments. Statistical testing was performed using a Mann-Whitney test.

### Optical trapping experiments

Experiments were performed on a custom-built, microscope free dual-beam optical trap described previously (29). These experiments utilized the three-bead geometry in which a thin filament is held between two optically trapped beads and lowered on to a surface bound bead that is sparsely coated with myosin (27,28). Tropomyosin was dialyzed into KMg25 buffer the night before the experiment. Actin was attached to beads using a biotin-streptavidin linkage, where actin contained 10% biotinylated actin and polystyrene beads were coated with streptavidin, as previously described (28,29). Flow cells were coated sparsely with beads as previously described (28,29). Flow cells were loaded with myosin (1-7 nM in KMg25 with 200 mM KCl to prevent myosin filament formation) for 5 minutes and the surface was blocked with 1 mg/mL BSA for 5 minutes. This was followed by activation buffer. For low ATP experiments, the activation buffer contained KMg25 with 1 μM ATP, 192 U/mL glucose oxidase, 48 μg/mL catalase, 1 mg/mL glucose, and ~25 pM biotin-rhodamine-phalloidin actin. For the high ATP experiments, conditions were identical, except 1 mM Mg*ATP was used. When appropriate, troponin and tropomyosin were also included at 200 nM. The concentration of free calcium was calculated using MaxChelator (60). This was followed by 4 μL of streptavidin beads. Flow cells were then sealed with vacuum grease as previously described and data were collected within 90 minutes of sealing (29).

Surface-bound beads were probed for binding interactions using small oscillations of the stage position that were stopped once data collection began. Data were collected at 20 kHz and filtered to 10 kHz according to the Nyquist criterion. For each bead-actin-bead assembly, the trap stiffness was calculated from fitting of the power spectrum as previously described (28).

### Implementation of the isometric optical clamp feedback

Here, we used an all-digital implementation of an isometric feedback clamp (36). In an isometric optical clamp feedback experiment, the position of one bead (the transducer) is continuously sampled, and deviations from its original setpoint position are compensated for by moving the second (motor) bead using acoustic optical deflectors (AODs, Gooch and Housego). The positions of the beads were recorded using quadrant photodiodes, the feedback calculations were digitally performed on a field programmable gate array (FPGA) board (National Instruments PCIe-7852), and the laser controlling the motor bead was translated using AODs.

The error signal used for the feedback, V_t_, is the time filtered positional error for the current sample period given by:

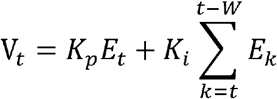

where K_p_ is the user defined proportional gain, K_i_ is the user defined integral gain, E_t_ is the current sample’s absolute error from the setpoint, and W is the user defined integration window (up to a 255-sample memory). The compensating position for the motor bead in frequency units is then calculated by the FPGA and transmitted to the parallel port interface of the Digital Frequency Synthesizer (Analog Devices AD9912A/PCBZ) which controls the beam deflection angle from the AOD. The feedback loop can run at a maximal speed of 50 kHz, limited by parallel port cable capacitance.

The time constant for the feedback response time was set as previously described (36). Briefly, a bead-actin-bead dumbbell was held in the dual beam traps, a square wave was injected into the transducer bead channel, and then movement of the motor bead by the feedback system was monitored. The proportional and integral gains were empirically adjusted to give a response time of ~5 ms without introducing oscillations into the system.

### Analysis of single molecule data

All data from optical trapping experiments were analyzed using our custom-built MATLAB program, SPASM, as previously described (29). Briefly, binding interactions between myosin and the thin filament were identified using a peak-to-peak covariance threshold, and the initiation and termination times of binding were identified using a changepoint algorithm. Data traces were excluded if the separation between the bound and unbound populations of the covariance histogram was not well defined. To improve the signal-to-noise ratio, the signal from both beads was summed and divided by 2 as previously described (29). Ensemble averages and histograms of binding interactions were generated as previously described (28–30). Ensemble averages were fit by single exponential functions in MATLAB until the signal plateaued. Cumulative distributions of step sizes and attachment durations were fit with cumulative functions for the Gaussian and exponential functions, respectively. Statistical testing for normality was done using a Shapiro-Wilk test. Statistical testing of the step sizes was done using a 2-tailed Student’s t-test of individual binding interactions. 95% confidence intervals for the detachment rate were calculated by bootstrapping of individual binding interaction durations, and statistical significance was calculated according to (62).

In the optical trap, actomyosin remains attached until ADP is released and ATP induces actomyosin dissociation (35):

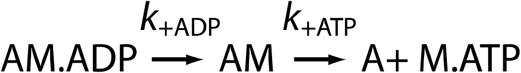

The attachment duration, t_on_, is given by

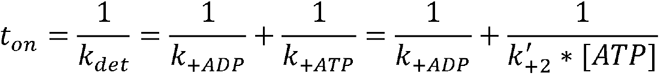

At low ATP, the detachment rate, k_det_, will be dominated by the second order rate of ATP binding, while at saturating ATP, the detachment rate will be limited by the rate of ADP release.

For isometric optical clamp experiments, data were collected at saturating ATP concentrations, where the rate of ADP release limits the rate of actomyosin dissociation (63). Binding interactions were identified using a variance threshold, set by the position of the transducer bead, and the force exerted by the motor bead and the attachment duration were measured. The relationship between the force, F, and the load dependent detachment rate, k(F) was modeled using the Bell equation (38):

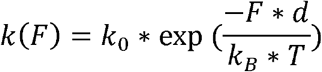

Where k_0_ is the rate of the primary force sensitive transition in the absence of force, d is the distance to the transition state, and k_B_*T is the thermal energy. The distribution of attachment durations is exponentially distributed at each force, and therefore follows the following probability density distribution (37):

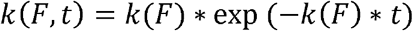

Maximum likelihood estimation (MLE) was used to determine the most likely values of k_0_ and d, as previously described (37). 95% confidence intervals for parameter values fitting were determined using 1000 rounds of bootstrapping simulations, and hypothesis testing was performed using the difference in the means and the variances of these distributions as previously described (62).

## Data Availability, Reproducibility, and Software

The data and code necessary to generate the figures and statistical tests in this manuscript are on Github (https://github.com/GreenbergLab/2023-thin-filament-trapping). Optical trapping data were analyzed with our previously published SPASM program by running the MATLAB source code from GitHub (https://github.com/GreenbergLab/SPASM). Similarly, the isometric force clamp data was analyzed with a modified version of SPASM (in the GitHub repository). Subsequent figure generation and related analyses were performed in R (version 4.2.2) (64) running under NixOS 22.05 (Linux). The git repository hosts a “Nix Flake”, a reproducible developmental shell environment with pinned version dependencies. For NixOS users, users of the Nix package manager on macOS or the Windows Subsystem for Linux (WSL), an ephemeral shell can be spawned by running *‘nix develop github:GreenbergLab/2023-thin-filament-trapping?dir=code’* in the terminal. Additional R packages used include the following: “readr_2.1.3”, “readxl_1.4.0”, “data.table_1.14.6”, “here_1.0.1”, “ggpubr_0.5.0”, “ggtext_0.1.2”, “purrr_0.3.5”, “tidyr_1.2.1”, “dplyr_1.0.10”, “tibble_3.1.8”, “cowplot_1.1.1”, and “ggplot2_3.4.0”.

## Funding Acknowledgements

This work was supported by the National Institutes of Health (R01 HL141086 to M.J.G.) and the Children’s Discovery Institute of Washington University and St. Louis Children’s Hospital (PM-LI-2019-829 to M.J.G.).

## Author contributions

S.R.C.S., W.T.S., B.S., and M.J.G. designed the experiments. S.R.C.S., B.S., and S.K.B. conducted and analyzed the optical trapping experiments. W.T.S. designed and tested the optical trapping system, including the feedback system. T.B., B.S., and M.J.G. contributed to tools for data analysis. S.R.C.S. drafted the first draft of the manuscript with M.J.G. All authors contributed to the analysis of data and writing/editing of the manuscript. M.J.G. procured funding and oversaw the project.

