## Supplemental Materials for "Single Molecule Mechanics and Kinetics of Cardiac Myosin Interacting with Regulated Thin Filaments" (from another version of this preprint: version 1, 2023-01-10)

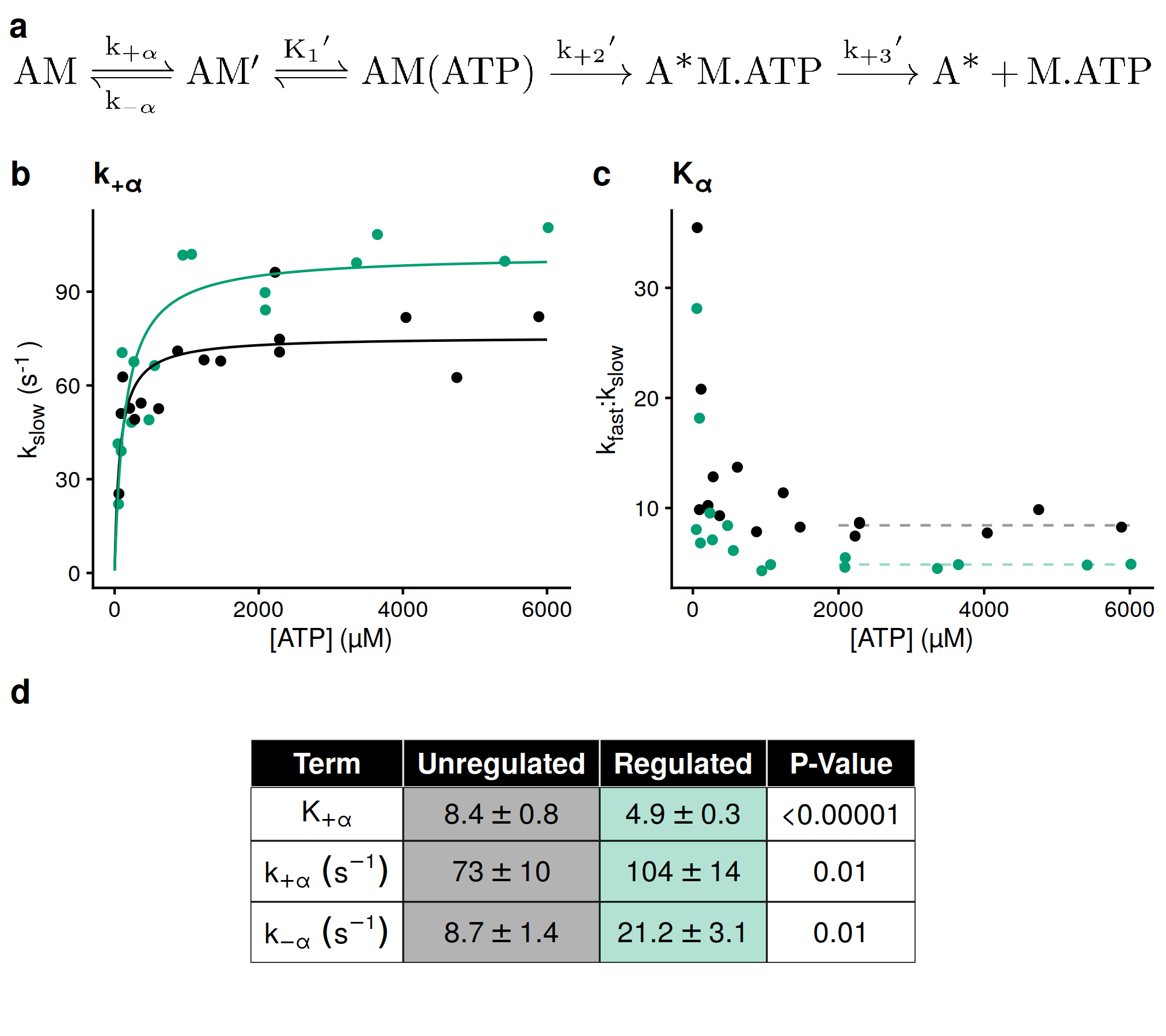

**Figure S1: Additional phases of ATP-induced dissociation measured in the stopped flow. a**. The kinetic scheme used to interpret data. **b.** Rate of the slow phase of ATP-induced actomyosin dissociation is plotted as a function of ATP. Regulated filaments are shown in green and unregulated are shown in black. Data were fitted with the following equation:

$\frac{\left( k_{+\alpha}\cdot\left[ ATP \right] \right)}{K + \left[ ATP \right]}$	Equation S1

where k_+α_ is the rate of the nucleotide-free isomerization (1,2). **c.** The ratio of the amplitudes of the fast to slow phases can be used to calculate the equilibrium constant, K_+α_. Regulated filaments are shown in green and unregulated are shown in black. K_+α,_ is obtained from the ratio of the amplitudes at high ATP concentrations (dashed lines). **d.** Table of parameters and values (mean +/- standard deviation) obtained from 3 separate experimental days. P-values are calculated by a 2-tailed Student’s t-test k_-α_ is calculated by dividing the rate of k_+α_ by K_+α,_and propagating the error.

1.	Geeves, M. A., C. Perreault-Micale, and L. M. Coluccio. 2000. Kinetic analyses of a truncated mammalian myosin I suggest a novel isomerization event preceding nucleotide binding. *J Biol Chem*. 275(28):21624-21630, doi: 10.1074/jbc.M000342200.

2.	Deacon, J. C., M. J. Bloemink, H. Rezavandi, M. A. Geeves, and L. A. Leinwand. 2012. Identification of functional differences between recombinant human alpha and beta cardiac myosin motors. *Cell Mol Life Sci*. 69(13):2261-2277, doi: 10.1007/s00018-012-0927-3.
